# Programmable Modulation for Extracellular Vesicles

**DOI:** 10.1101/566448

**Authors:** Yihua Wang, Richard Melvin, Lynne T. Bemis, Gregory A. Worrell, Hai-Long Wang

## Abstract

Every living cell releases extracellular vesicles (EVs) that are critical for cellular signaling and a wide range of biological functions. The potential diagnostic and therapeutic applications of EVs are well recognized, and rapidly expanding. While a complete understanding of the molecular mechanisms underpinning EVs release remains elusive, here we demonstrate a novel method for programmable control of the release of EVs and their cargo using external electric fields. As a proof of principle, we use cultured rat astrocytes to demonstrate how the frequency of external electrical stimulation selectively modulates EV release, their surface proteins, and microRNA profiles. This method could broadly impact biological science and medical applications. First, it raises an interesting question of how endogenous electrical activity could modulate EV production. Second, it provides a novel mechanism for tuning therapeutic electrical stimulation that may be useful for treating brain disorders. Third, it provides a new way to generate EVs carrying desired cargos by tuning electrical stimulation parameters. Unlike chemical methods for creating EVs, electrical stimulation is a clean physical method with adjustable parameters including stimulation frequency, field strength and waveform morphology.

---

EVs are membrane-enclosed structures released by cells in an evolutionally conserved manner. All cell types tested to date produce EVs. They are widely distributed in blood, cerebral spinal and other bodily fluids. These novel signaling mediators carry bioactive molecules between cells (*1*), and studies suggest they have both physiological and potentially pathological roles in viral infection (*2*), cancer spread (*3*), autoimmunity (*4*) and neurodegenerative diseases (*5*). EVs transfer their protein, lipid and RNA cargo in both paracrine (to adjacent target cells) and endocrine manner (to distant target cells) (*6*). Because of these unique properties, EV-related diagnostic and therapeutic applications are under intense investigation.

The production and release of EVs is a tightly regulated process, varying between physiological and pathologic conditions (*7, 8*). External stimuli can profoundly change the production rate and content or composition of EVs. However, chemical stimulation lacks specificity and releases a heterogeneous population of vesicles (*9, 10*). Ideal therapeutic EV production requires clean and efficient regulation.

The mammalian brain generates endogenous continuous electric field activity with extracellular voltages less than 0.5 mV and fields under 5 mV/mm (*11*–*14*). External electrical stimulation, using invasive or non-invasive electrodes to target selected brain regions, is an emerging neuromodulation therapy. Fields ranging from 0.5mV/mm ~ 100mV/mm are able to forestall or even reverse nervous system dysfunction, and focal external electrical stimulation is an FDA-approved treatment for multiple neurological diseases (*15*). The prevailing theory for its mechanism of action is that neuronal activity can be generated or blocked by extracellular electrical stimulation, and that different frequencies selectively affect different neuronal pathways (*16*). Interestingly, many electrical stimulation therapies report improved efficacy over long time-scales, e.g. months to years (*17, 18*). The mechanisms by which electrical stimulation remodels the nervous system are not well understood. The study that we now report suggests that EVs could be mediators of therapeutic external electrical stimulation. The application of programmable external electrical stimulation to control the release of EVs and their cargos is a generic concept applicable to many cell types.

As a proof of principle, we chose to study astrocytes, the most abundant (4-5 fold more numerous than neurons) and heterogeneous of glia in the central nervous system (CNS). Their synaptic interactions with neuronal synapses and nodes of Ranvier, blood vessels and other glial cell types (*19*–*21*) situates them optimally for modulating neurons. Known functions (*22*) include regulation of synaptic connectivity, integration and synchronization of neuronal networks and maintenance of the blood–brain barrier integrity. They respond to low levels of electrical stimulation by elevation of their intracellular calcium concentration (*23, 24*). In our initial testing, we found that ambient electrical stimulation affects EV release from rat astrocytes in primary culture (Figure 1). We used a green fluorescent dye, Calcein, to monitor cytoplasmic volume and vesicle extrusion activity of astrocytes (see supplement 1 for method and materials). Outward budding and vesicle shedding was observed at the plasma membrane during electrical stimulation (2 Hz square wave with 0.1 ms pulse width). Figure 1C shows image of EVs released during electrical stimulation. The applied field magnitude was determined by trial stepwise lowering of the field strength from 100mV/mm. We found that most cells were killed at high field strength, but at or below ~5 mV/mm most cells survived electrical stimulation. This field magnitude is within the range of endogenous electrical fields generated by neurons in the hippocampus (*14*).

**Figure 1.**
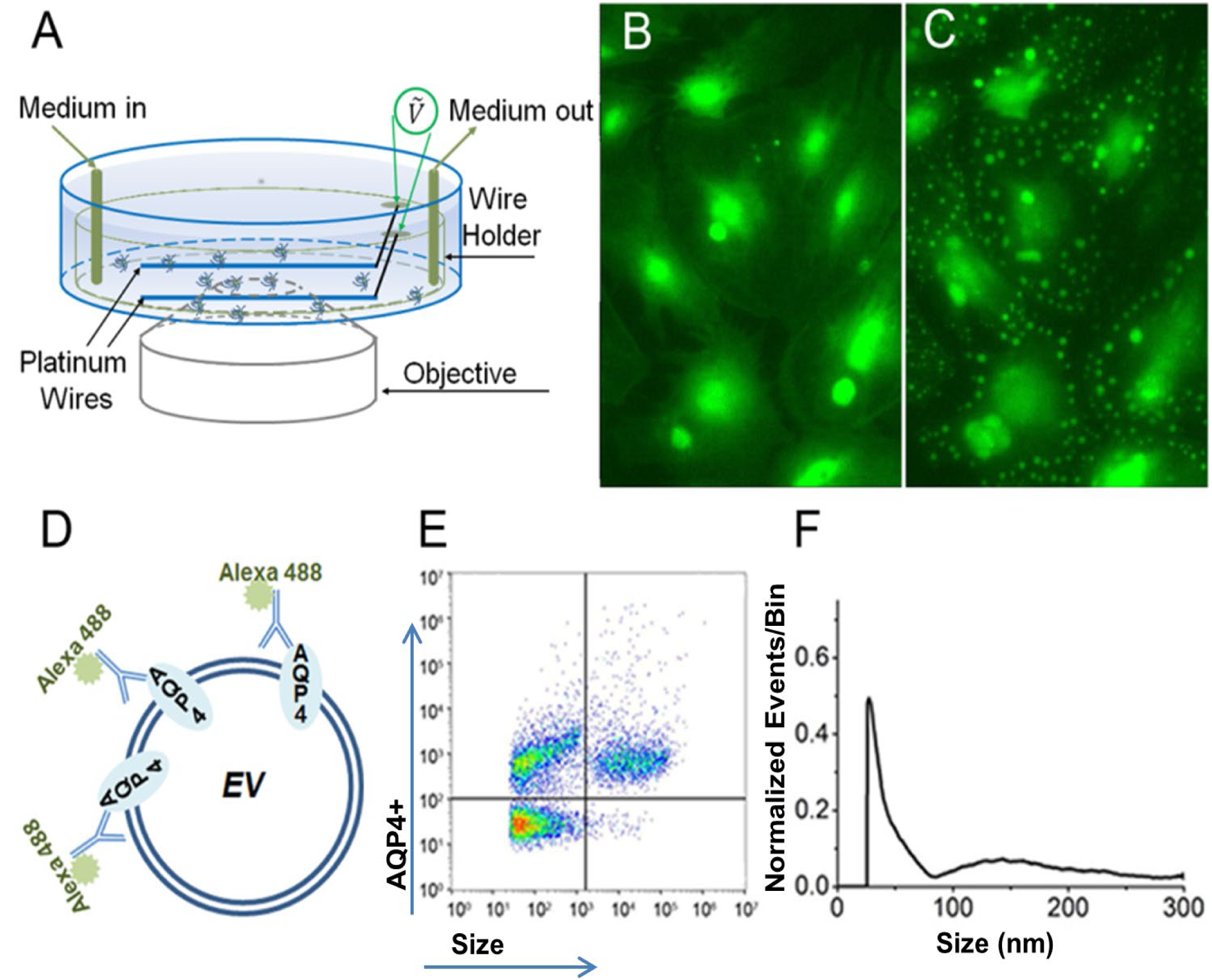
Extracellular vesicles (EVs) budding off the plasma membrane of astrocytes and EV detection with nano-flow cytometry. **1A** Schematic of electrical stimulation chamber for culture astrocytes plated in a petri dish. Cultured astrocytes loaded with Calcein (a green fluorescent dye) are shown before (**1B**) and after (**1C**) electrical stimulation (2 Hz square wave with 0.1 ms pulse width and 5mV/mm field). Release of EVs through budding off the plasma membrane from astrocytes was observed using an high-magnification object lens. **1D** Drawing of EV with membrane AQP4. Alexa Flour 488 fluorescence conjugated monoclonal antibody IgG to the extracellular domain of AQP4 is used for EV labeling. **1E** & **1F** show a typical bivariate dot-plot of fluorescence (FITC) versus side scattering (SSC) and the corresponding size distribution of EVs.

To further determine the selective effect of electrical stimulation waveforms on the release of astrocytic EVs, we applied the same strength (5 mV/mm) electric field at three different frequencies that have been used in therapeutic electrical brain stimulation in humans, 2 Hz, 20 Hz and 200 Hz. In fact, these frequencies also fall into three endogenous brainwave oscillation frequencies recorded with human intracranial electroencephalography (iEEG), delta (0.5 - 4 Hz), beta-gamma (>12 - 30 Hz) and ripple (100 - 200 Hz). Here we applied electrical stimulation to astrocyte cultures through a custom rig fitted to a 100 mm petri dish (Figure 1A) designed to create uniform electric fields over the astrocytes with continuous fluid exchange to collect EVs.

To characterize EVs in detail, we purified collected EVs using ultrafiltration combined with size-exclusion chromatography to separate EVs from extrinsic proteins (*25*), and then used a nano-flow cytometer (N30 Nanoflow Analyzer, NanoFCM Inc., Xiamen, China) to quantify EVs by their size and surface proteins (see supplement 1 for method and materials). In this case, we monitored the astrocyte-specific water channel, aquaporin-4 (AQP4), by using an Alexa Flour 488 conjugated monoclonal AQP4 extracellular domain-specific IgG (Figure 1D). We used AQP4 as an indicator for particle numbers and sizes corresponding to astrocytic EVs collected at different stimulation frequencies. In control condition without applied electrical stimulation, the nano-flow cytometry plot shows populations of AQP4-positive EVs of two distinct sizes (Figure 1E).

In general, EVs are classified into three major groups: exosomes, microvesicles and apoptotic bodies based on vesicle diameter and the release process (*26*). Exosomes are the smallest vesicles (~ 50 - 100 nm) released through exocytosis of endosomal multivesicular bodies. Microvesicles, also known as ectosomes (~100 - 1000 nm), are generated by outward budding and shedding from the plasma membrane. Apoptotic bodies (> 1000 nm) are released from apoptotic cells. We interpreted the smaller sized vesicles to be exosomes and the larger sized vesicles to be microvesicles. Thus, we concluded that the vesicles we visualized arising from cultured astrocytes under electrical stimulation in Figure 1C were most likely microvesicles, given the limited resolution of optical microscopy (> 100 nm).

In this experiment, we found that the size distributions of AQP4-positive EVs are differentially affected by the frequency of electrical stimulation (Figure 2). Stimulation at 2 Hz produces a near uniform distribution of EVs, with the peak of size distribution at ~70 nm, slightly shifted to the right (larger) by comparison with the smaller sized presumptive exosome population collected without electrical stimulation. The major population of AQP4-positive exosomes arising from 2 Hz stimulation was more AQP4-bright than at baseline, supporting that 2 Hz stimulation generated a population of exosomes with increased membrane AQP4. Stimulation at 20 Hz produced fewer EVs of both large and small size than seen in basal conditions, particularly fewer presumptive exosomes. Electrical stimulation at 200 Hz had the opposite effect of 2Hz stimulation. Instead of yielding exosomes, microvesicles-sized EVs dominated (peak size distribution at ~ 170nm) and this population was also enriched in membrane AQP4.

**Figure 2.**
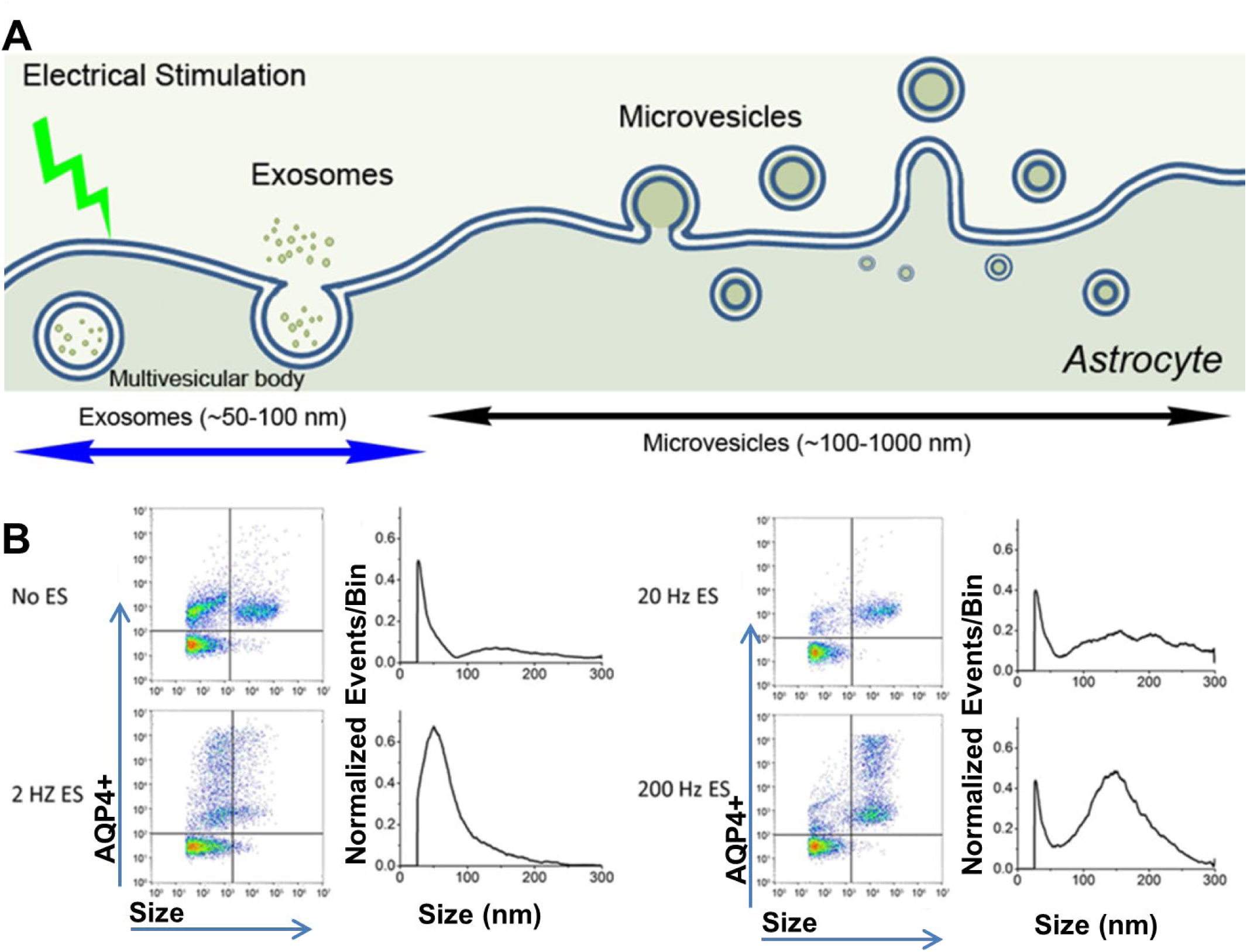
Effect of electrical stimulation on EV cargos and size distributions. **2A** illustrates our hypothesis that the frequency of an electrical field applied to astrocytes affects both exosomes and microvesicles. **2B** Shows bivariate dot-plots and size distributions of AQP4^+^ EVs analyzed by nano-flow cytometry. EVs were collected without electrical stimulation, at 2 Hz stimulation, at 20 Hz and at 200 Hz.

The characteristics of the evoked EVs demonstrate a complex range of variability that not only depends on the parameters of stimulation, but also on the preparation of the astrocyte cultures. We tested three independent batches of cultured astrocytes prepared under slightly different protocols. Despite some variability in preparation and the response to ES, we investigated the percent change in exosome and microvesicle release (see supplementary 2 for additional data). The microvesicles increased under electrical stimulation (p<0.05), but the exosomes show a more variable effect at different frequencies. This variability in response to external electrical stimulation from cells prepared under different conditions suggests the potential for future precise treatment using electrical stimulation (see more in later discussion).

The RNA cargo is an important property of EVs. Stress of many types is known to increase the small noncoding RNA contained in EVs (*27*). We therefore anticipated that the cellular response of astrocytes to low amplitude electrical stimulation might affect miRNA cargos of EVs. We took an unbiased approach to investigate the effect of electrical stimulation frequency on miRNA profiles by using RNA-seq to identify any changes in EV miRNAs. We extracted RNAs from purified EVs collected respectively after applying a uniform electric field (5 mV/mm) at 2 Hz, 20 Hz, and 200 Hz electrical stimulation, using reverse transcription to convert them to cDNA library for high-throughput sequencing. RNA-seq results are summarized in a Venn diagram (Figure 3) and details are listed in Table 1. We identified a total of 89 miRNAs. The number of miRNAs identified in each sample of EVs collected under four conditions were: No electrical stimulation, 32 miRNAs; 2Hz stimulation, 45 miRNAs; 20Hz stimulation, 65 miRNAs and 200Hz stimulation, 67 miRNAs. Twenty one miRNAs were present in all four groups, representing 66% in the control group with no electrical stimulation, 47% with 2Hz, 32% with 20Hz and 31% with 200 Hz. Altogether, fourteen miRNAs were specifically associated with electrical stimulation regardless of stimulation frequency. However, each of the three electrical stimulation frequencies studied had a unique set of miRNAs.

**Table 1.**
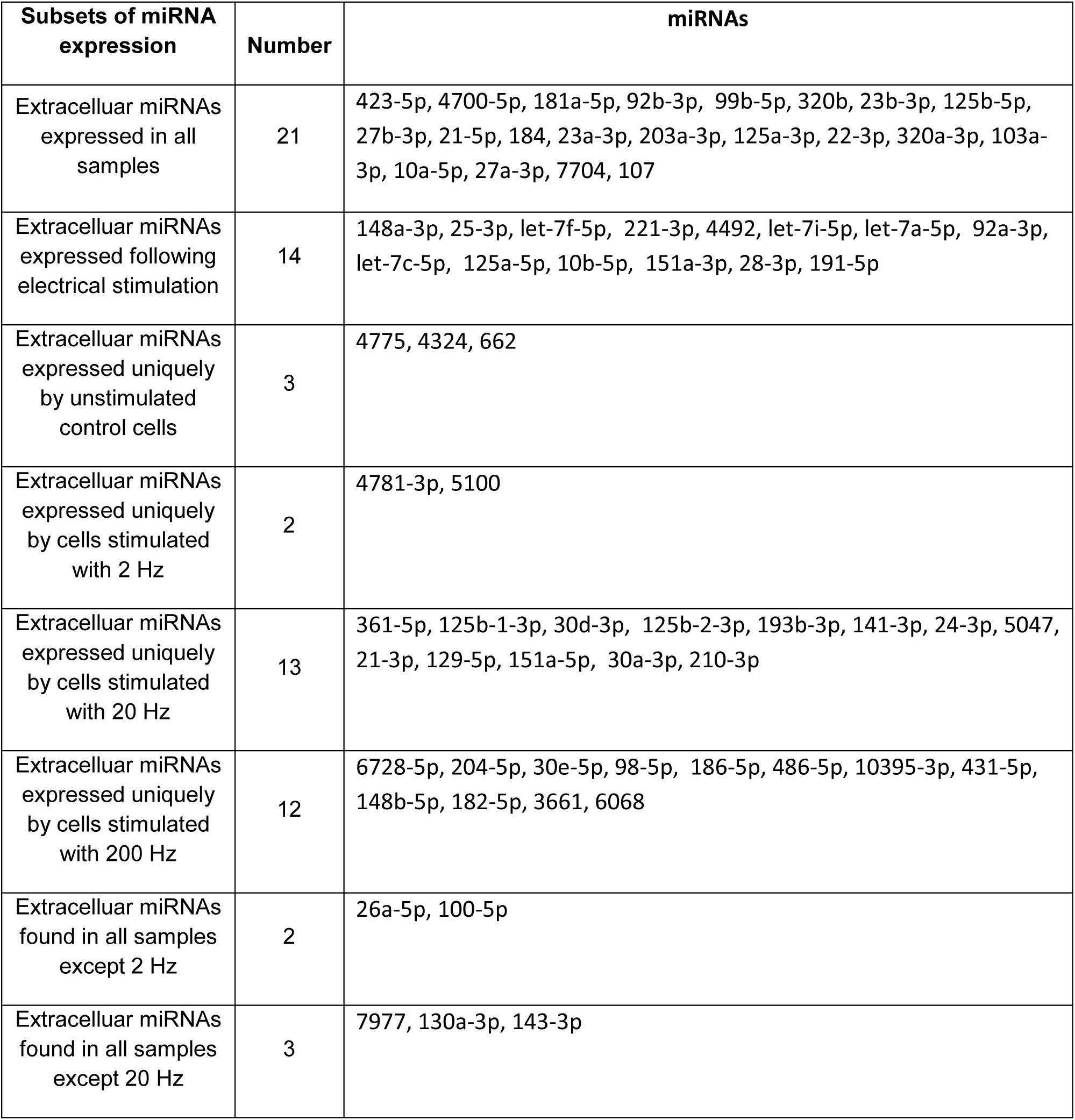
MiRNAs distributions

**Figure 3.**
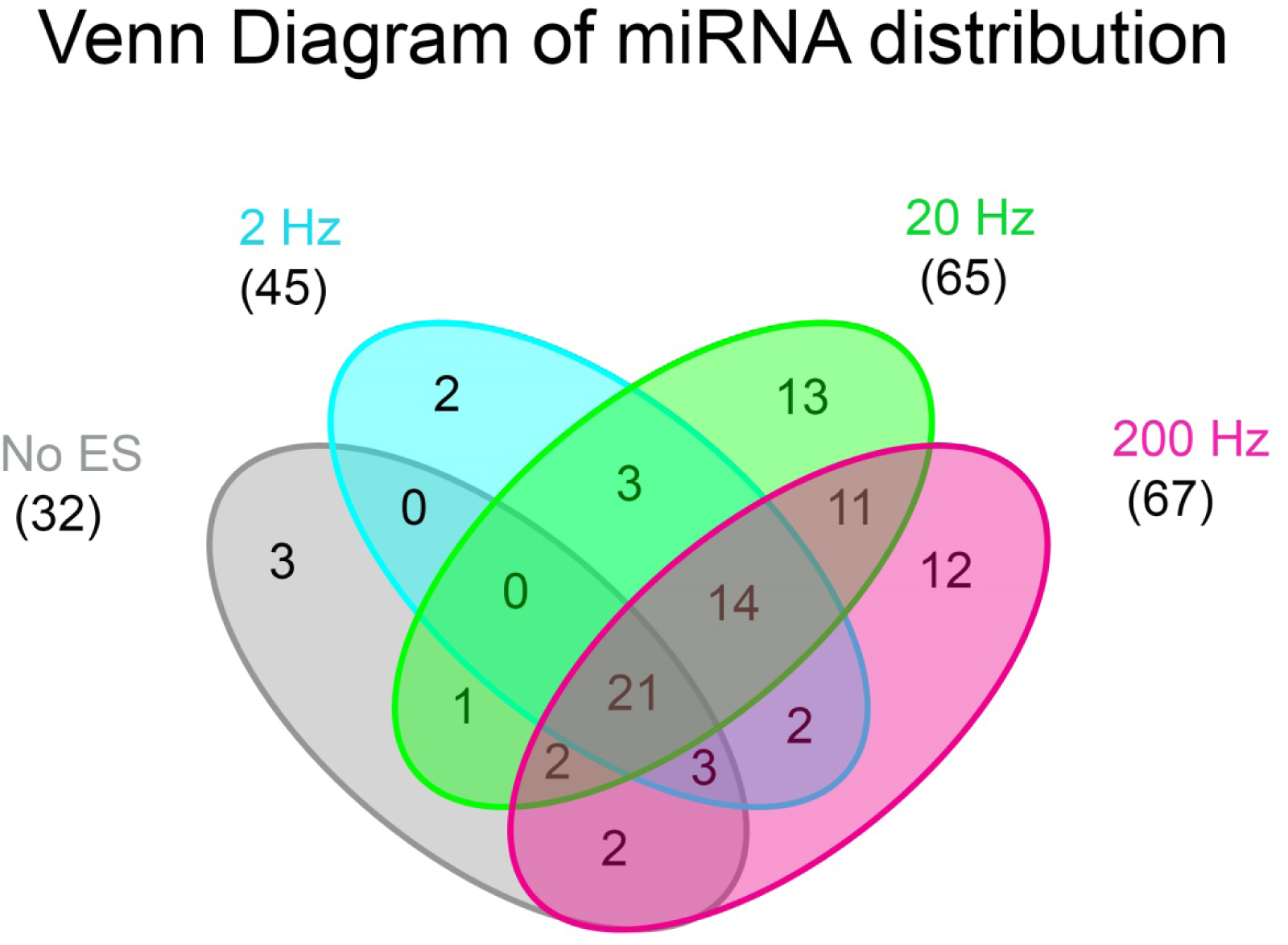
A Venn diagram of miRNA distribution from RNA-seq.

In summary, we have demonstrated that the number, size, surface proteins, and cargo of EVs released from cultured rat astrocytes is influenced selectively by the frequency of electrical stimulation. Perhaps not surprisingly, the characteristics of EV release from cultured astrocytes at baseline (no electrical stimulation) and under different electrical stimulation conditions depend on astrocyte culture preparation. We have successfully tested three batches of culture astrocytes prepared under slightly different protocols. Despite the limited range of experimental data and certain variabilities in terms of EV release, our general conclusion is that electrical stimulation can selectively modulate release of EVs. In fact, variable effect of electrical stimulation on cells under different physiological conditions suggests that normal and diseased cells will respond differently to electrical stimulations. Future methods may be designed to target only diseased cells and have less or even no effects on normal cells.

In the future, we also need to examine effects of electrical stimulation on lipids and other astrocytes-specific proteins, such as the glutamate transporter and the inward-rectified potassium channel. However, the key point emerging from this initial study is that electrical stimulation provides a clean physical method for programmable modulation of EVs and their cargo that could have a broad impact in biological science and medical applications.

The observations we report also raise interesting questions concerning the mechanism by which external and endogenous electrical events modulate EV release. First, the observed effect may represent a fundamental phenomenon of possible pertinence to the response of primitive single-cell life-forms to external electrical fields. This may be important for the possible biological role of large-scale electrical activity of the mammalian brain. The electroencephalogram (EEG) reveals several frequency bands in the brain’s endogenous electrical activity: delta (1–4 Hz), theta (>4–8 Hz), alpha (>8– 13 Hz), beta (>13–30 Hz), low gamma (>30–70 Hz), high gamma (>70–150 Hz) and fast ripples (>250-600 Hz). It is thought that local field potentials modulate the spike timing of individual neurons via ephaptic coupling (*11*). One of the most dramatic changes observed in the mammalian EEG occurs during transition from the wake state to sleep where the EEG activity changes to higher amplitude and low frequency. The coinciding changes in electrical activity and behavioral state may shape the profile of released EVs and their cargo composition. Our study suggests that endogenous extracellular electrical activity has a potent potential to modulate the release and cargo composition of EVs to serve various biological and pathological functions.

Second, our results suggest a means for biologically monitoring the tuning of therapeutic electrical stimulation in the treatment of several neurological disorders (*28, 29*), such as epilepsy (*17, 30*–*32*), sleep and memory dysfunction (*33*–*35*), brain tumors (*36*) and Parkinson disease (*37*). It has been noted empirically that the frequency of electrical stimulation is a critical determinant of outcome in treating particular neurological diseases. For parkinsonian and idiopathic tremor, high frequencies (>90Hz) improve motor symptoms, but lower frequencies (<50Hz) are either ineffective or can exacerbate symptoms (*38*). By contrast, in some epilepsy studies low frequency stimulation (< 3Hz) safely and effectively reduces seizures (*30, 39*). The mechanism underlying success of frequency-dependent electrical stimulation therapy is still largely unknown. Monitoring of EV profiles in brain cerebrospinal fluid, serum, or interstitial fluid (by dialysis) may provide measurable biomarkers for optimizing electrical stimulation therapy to achieve a better clinical outcomes.

Third, the results suggest a new method to produce therapeutic EVs carrying desired cargos by tuning electrical stimulation parameters *in vitro*. Unlike chemical methods for creating EVs, electrical stimulation is a clean physical method with several adjustable parameters including oscillation frequency, field strength and waveform.

## Acknowledgements

This study was funding by the Minnesota Partnership for Biotechnology and Medical Genomics to L.B., G.W. (MNP #17.16). We are grateful to Drs. Ling Ma and ShaoBin Zhu from nanoFCM who provided us the Nano Analyzer as loan equipment and technical assistance on preparing samples. We thank Dr. Vanda A Lennon for providing us astrocyte cultures and monoclonal AQP4 antibody IgG. We also thank Ms. Ningling Luo and Jacquelyn A. Grell who prepare astrocyte cultures.

## Author Contributions

H. W. designed experiments; H.W., Y.W., L.B. and R.M. carried out the experiments and analysis; H.W., G.W. L.B. wrote the paper.

## Competing financial interests

The authors declare no competing financial interests

Table 1. miRNAs identified from RNA-seq.

## References

1. K. W. Witwer et al., Standardization of sample collection, isolation and analysis methods in extracellular vesicle research. J. Extracell. Vesicles. 2, 20360 (2013).

2. M. Rodrigues, J. Fan, C. Lyon, M. Wan, Y. Hu, Role of Extracellular Vesicles in Viral and Bacterial Infections: Pathogenesis, Diagnostics, and Therapeutics. Theranostics. 8, 2709–2721 (2018).

3. A. Becker et al., Extracellular Vesicles in Cancer: Cell-to-Cell Mediators of Metastasis. Cancer Cell. 30 (2016), pp. 836–848.

4. D. Turpin et al., Role of extracellular vesicles in autoimmune diseases. Autoimmun. Rev. 15, 174–183 (2016).

5. T. Croese, R. Furlan, Extracellular vesicles in neurodegenerative diseases. Mol. Aspects Med.60, 52–61 (2018).

6. X. Yu, M. Odenthal, J. W. U. Fries, Exosomes as miRNA carriers: Formation-function-future. Int. J. Mol. Sci.17 (2016), doi:10.3390/ijms17122028.

7. M. Yáñez-Mó et al., Biological properties of extracellular vesicles and their physiological functions. J. Extracell. Vesicles. 4 (2015), pp. 1–60.

8. J. De Toro, L. Herschlik, C. Waldner, C. Mongini, Emerging roles of exosomes in normal and pathological conditions: New insights for diagnosis and therapeutic applications. Front. Immunol.6, 1–12 (2015).

9. E. N.M. Nolte-’t Hoen, M. H.M. Wauben, Immune Cell-derived Vesicles: Modulators and Mediators of Inflammation. Curr. Pharm. Des.18, 2357–2368 (2012).

10. E. J. van der Vlist et al., CD4+ T cell activation promotes the differential release of distinct populations of nanosized vesicles. J. Extracell. Vesicles. 1, 1–9 (2012).

11. C. A. Anastassiou, R. Perin, H. Markram, C. Koch, Ephaptic coupling of cortical neurons. Nat. Neurosci.14, 217–224 (2011).

12. G. Buzsáki, Z. Horváth, R. Urioste, J. Hetke, K. Wise, High-frequency network oscillation in the hippocampus. Science (80-.).256, 1025–1027 (1992).

13. A. Kamondi, L. Acsády, X. J. Wang, G. Buzsáki, Theta oscillations in somata and dendrites of hippocampal pyramidal cells in vivo: Activity-dependent phase-precession of action potentials. Hippocampus. 8, 244–261 (1998).

14. S. Weiss, D. Faber, Field effects in the CNS play functional roles. Front. Neural Circuits. 4, 1–10 (2010).

15. W. Deeb et al., Proceedings of the Fourth Annual Deep Brain Stimulation Think Tank: A Review of Emerging Issues and Technologies. Front. Integr. Neurosci.10, 38 (2016).

16. F. Rattay, The basic mechanism for the electrical stimulation of the nervous system. Neuroscience. 89, 335–346 (1999).

17. G. K. Bergey et al., Long-term treatment with responsive brain stimulation in adults with refractory partial seizures. Neurology. 84, 810–817 (2015).

18. V. Salanova et al., Long-term efficacy and safety of thalamic stimulation for drug-resistant partial epilepsy. Neurology. 84, 1017–25 (2015).

19. R. D. Fields, B. Stevens-graham, New Insights into Neuron-Glia Communication. Science (80-.).298, 556–562 (2002).

20. G. Perea, M. Sur, A. Araque, Neuron-glia networks: integral gear of brain function. Front. Cell. Neurosci.8, 378 (2014).

21. S. Hinson et al., Pathogenic potential of IgG binding to water channel extracellular domain in neuromyelitis optica. Neurology. 69, 2221–2231 (2007).

22. M. Nedergaard, B. Ransom, S. a. Goldman, New roles for astrocytes: Redefining the functional architecture of the brain. Trends Neurosci.26, 523–530 (2003).

23. H. Monai, H. Hirase, Astrocytic calcium activation in a mouse model of tDCS—Extended discussion. Neurogenesis. 3, 1–7 (2016).

24. H. Monai, H. Hirase, Astrocytes as a target of transcranial direct current stimulation (tDCS) to treat depression. Neurosci. Res. 126 (2018), pp. 15–21.

25. J. Z. Nordin et al., Ultrafiltration with size-exclusion liquid chromatography for high yield isolation of extracellular vesicles preserving intact biophysical and functional properties. Nanomedicine Nanotechnology, Biol. Med.11, 879–883 (2015).

26. M. Piffoux et al., Extracellular vesicles for personalized medicine: The input of physically triggered production, loading and theranostic properties. Adv. Drug Deliv. Rev.(2018), doi:10.1016/j.addr.2018.12.009.

27. K. W. Diebel, K. Zhou, A. B. Clarke, L. T. Bemis, Beyond the ribosome: Extra-translational functions of tRNA fragments. Biomark. Insights. 11, 1–8 (2016).

28. H. S. Mayberg et al., Deep brain stimulation for treatment-resistant depression. Neuron. 45, 651–660 (2005).

29. M. A. Nitsche et al., Transcranial direct current stimulation: State of the art 2008. Brain Stimul.1 (2008), pp. 206–223.

30. K. B. Kile, N. Tian, D. M. Durand, Low Frequency Stimulation Decreases Seizure Activity in a Mutation Model of Epilepsy. Epilepsia. 51, 1745–1753 (2010).

31. R. S. Fisher, Therapeutic devices for epilepsy. Ann. Neurol.71, 157–168 (2012).

32. S. Toprani, D. M. Durand, Long-lasting hyperpolarization underlies seizure reduction by low frequency deep brain electrical stimulation. J. Physiol.591, 5765–5790 (2013).

33. M. Z. Koubeissi, F. Bartolomei, A. Beltagy, F. Picard, “Electrical stimulation of a small brain area reversibly disrupts consciousness” (2014), doi:10.1016/j.yebeh.2014.05.027.

34. Y. Ezzyat et al., Direct Brain Stimulation Modulates Encoding States and Memory Performance in Humans. Curr. Biol.27, 1251–1258 (2017).

35. M. T. Kucewicz et al., Evidence for verbal memory enhancement with electrical brain stimulation in the lateral temporal cortex. Brain. 141, 971–978 (2018).

36. R. Stupp et al., NovoTTF-100A versus physician’s choice chemotherapy in recurrent glioblastoma: a randomised phase III trial of a novel treatment modality. Eur. J. Cancer. 48, 2192–202 (2012).

37. G. Deuschl et al., A Randomized Trial of Deep-Brain Stimulation for Parkinson’s Disease. N. Engl. J. Med.355, 896–908 (2006).

38. L. Wojtecki et al., The treatment of Parkinson’s disease with deep brain stimulation: current issues. NEURAL Regen. Res.10 (2015), doi:0.4103/1673-5374.160094.

39. B. N. Lundstrom et al., Chronic subthreshold cortical stimulation to treat focal epilepsy. JAMA Neurol.73 (2016), pp. 1370–1372.

