## Supplementary material for "Programmable Modulation for Extracellular Vesicles": Methods and Supplemental Data

### Materials and Methods

#### Astrocyte Cultures.

Pregnant Lewis rats from Harlan Sprague–Dawley were purchased from Envigo Labs. Animal experiments were performed under Mayo Clinic IACUC-approved protocol (A66014) and complied with NIH guidelines. Glial cultures, established from embryonic days 18–21 fetal rat cerebral cortices using McCarthy de Vellis method (1), were used at weeks 2–5. Microglia depletion is achieved by adding Clodrosome (100 µg/ml; Encapsula Nano Sciences LLC) to growth medium (DMEM, 10% FCS, sodium pyruvate, glutamine, antibiotics) for 12 h, with fresh medium substituted for at least 48 h, and was confirmed by Iba-1 immunostaining (Wako-chem. Osaka, Japan). For fluorescent imaging, subculture bulk astrocytes to 35mm dishes with No. 1.0 glass coverslip pre-coated with Poly-D-lysine (P35GC-1.0-14-C from MatTek. Ashland, MA) and with additional coating of laminin. Cell density was about $2.5x10^5$cell count. Hold at $37^oC/5\%CO_2$/humid air for 48 hours. For EV collection, subculture bulk astrocytes to 100 mm Corning™ Falcon™ Bacteriological petri dishes (#351029 Corning, Corning NY) coated with Poly-D-lysine before use. Cell density between 1~5 $x10^5$ count.

#### Cell Imaging

We used an Olympus inverted fluorescence microscope IX73 to image cells at high magnification. Cells grown on glass coverslips in 35 mmm MatTek dishes were loaded with 10 uM calcein, AM (Lot# 1853976 from Invitrogen by Thermo Fisher Scientific. Waltham, MA) and maintained at 37 $^oC$ for 40 mins before imaging. The cells were illuminated with Xenon Arc bulb housed in a Lambda LS (Sutter Instrument. Novato, CA) with a filter set for GFP (ex488/em520) imaged with a 25X dry objective. Emitted fluorescence was detected by a 16-bit interline CCD camera (Andor Clara, Oxford Instrument. Belfast, Northern Ireland) recorded at 3 fps with a smart shutter controller (Lambda 10-3, Sutter Instrument) using MetaMorph Advance software included in the package of MetaMorph for Olympus. ImageJ was used for later image processing. During imaging acquisitions, cells were maintained at 37 $^oC$ using a culture dish heater DH-35 (Warner Instruments) connected to a heater controller TC-324C (Warner Instruments).

#### Electrical Field Stimulation.

Cell culture media was aspirated three times with pre-heated 0.2 µm filtered isotonic PBS. Electrical field stimulation was applied through a dish insert with two platinum wires embedded in parallel to provide an evenly distributed electrical field within the open area. For 35mm dishes we used an insert (RC-37FS from Warner Instruments. Hamden, CT). Distance between two platinum wires in RC-37FS is 6 mm; for 100mm dishes, we used a custom-made insert, similar to RC-37FS but in a larger scale. The distance between two platinum wires is 24 mm. We used a stimulus isolator (A365, World Precision Instruments. Sarasota, FL) with a 1KΩ bridging resistor to output voltages. For 35 mm dishes, setting on A365 is 100 µA with a ratio at 30% that generates a voltage of 30 mV (final field strength is 5mV/mm). For 100 mm dishes, setting on A365 is 1000 µA with a ratio at 12% that generates a voltage of 120mV (final field strength is still 5mV/mm). Stimulation parameter is controlled by the software (PClamp 10) that generates square waves with 0.1 millisecond in pulse width on a Digidata 1440 (Molecular Device. San Jose, CA) that triggers the stimulus isolator A365. We added resting times between clusters of pulses so that final number of applied pulses is always the same regardless of the simulation frequency. As the result, a longer resting time was used for stimulation at a higher stimulation frequency.

#### EV purifications for EV surface protein analysis

EVs were collected from cells grown in 100 mm dishes under 37 °C with a custom-made dish heater similar to the DH-35. We aspirated cell culture media three times with pre-heated 0.2 μm filtered isotonic PBS before EV collection. After 5 min of electrical stimulation, 10 ml of PBS media were collected using a custom-made automated solution exchange system. Collected PBS media was first filtered with 0.22 μm (Millipore. Burlington, MA), and then centrifuged in conical tubes at 3,000 x g for 20 mins at 4 °C to remove cell debris. 500 μl of concentrated EV media was achieved using Amicon Ultra-15 Centrifugal Filter Units with 10K cutoff (UFC901014, Millipore. Burlington, MA) spin at 4,000 x g for 13 mins at 4 °C. Further EV purification was done through a size-exclusion column (qEVoriginal from Izon, Medford, MA).

**Nano-flow cytometry**

Nano-flow cytometry analysis was performed as described (2) with minor modifications. Basically, 250 μl of purified EV samples were further concentrated using ultracentrifugation spin at 100,000 x g (TA100.3) at 4°C for 45 mins. Pellets were re-suspended in 50 μl of vesicle-free PBS, and then incubate with 2μl of Alexa Fluor 488 (A10235, Life Technologies, Grand Island, NY) conjugated monoclonal AQP4 antibody (specific to AQP4 external domain) at 37 °C for 30 mins. The mixture was washed twice with PBS by ultracentrifugation at 100,000 × g (TLA100 rotor) at 4 °C for 20 min. The pellet was re-suspended in 25 μl of PBS for Nano-flow cytometry analysis using the Nano Analyzer N30 (nanoFCM, Fujian, China).

**Total EV and extracellular RNA collection**

PBS was collected and frozen to allow for total extracellular RNA isolation. All subsets of EVs and extracellular proteins that might be associated with extracellular RNA are collected in this manner. Samples were then sent on ice to the UMN campus in Duluth for RNA analysis. Upon arrival at UMN campus all samples were immediately aliquoted into 50 μl aliquots with 700 μl s of Qiazol (Qiagen, Valencia, CA) added to each aliquot. All samples were then stored at -80°C until RNA extraction.

***RNA extraction and precipitation***

RNA extraction was conducted essentially as previously described (3). Briefly, total RNA was extracted from each of the five 50 μl aliquots using the miRNeasy Mini Kit (Qiagen). RNA was removed from the MiRNeasy spin column with 40 μl of Rnase free water and all five aliquots were combined for precipitation (200 μl). Total RNA was precipitated using 1 μl of GenElute LPA (Sigma-Aldrich, St. Louis, MO), 3 mM NaAcetate and ice cold 100% ethanol. The use of GenElute is necessary because alternatives derived from natural sources, such as glycogen may harbor small RNAs that would be sequenced when the samples are subjected to Next Gen Sequencing. RNA was re-suspended in 15 μl of water and the concentration and quality were assessed by spectrophotometry on the NanoDrop 100 (Thermo Scientific, Waltham, MA).

***RNA amplification***

The precipitated RNA was then used in two protocols. First, 2 μl of RNA was subjected to cDNA preparation using the miScript II RT kit (Qiagen) for reverse transcription. These samples were then analyzed by qRT-PCR using the miScript SYBR Green reagent (Qiagen) with a custom primer for miR-21-5p 5'- ccc TAG CTT ATC AGA CTG ATG TTG A to confirm that amplifiable RNA was obtained as previously described (4). The remainder of the RNA was used in the Next Gen protocol successfully by ignoring the suggested quantity of RNA.

***Library preparation and Illumina Sequencing***

Small RNA libraries were constructed using Tru-Seq Small RNA Library Prep Kit -Set A or Set B (Illumina, San Diego, CA) according to the manufacturer's instructions. Each sample was barcoded and one set of the triplicate samples was included in each of three libraries. Small RNA libraries were then sequenced using next-generation sequencing on the Illumina HiSeq 2500 using high-output mode with paired-end sequencing with 50 cycles and v4 chemistry.  Prior to sequencing of the libraries, the libraries passed quantification testing using PicoGreen DNA quantification and Agilent Analysis for DNA. All Next Generation Sequencing and library testing was conducted at the University of Minnesota Genomics Center (UMGC), Minneapolis, MN.

NCBI Submission ID is SUB5033116; BioSample accession ID is SAMN10721961; BioProject ID is PRJNA514311. Project information will be accessible with the following link, http://www.ncbi.nlm.nih.gov/bioproject/514311.

***RNA Bioinformatics analysis***

Sequencing data was quality checked using FastQC (Babraham Bionformatics, Camebridge, UK) and trimmed of sequencing adapters using CutAdapt (5). The trimmed reads were mapped to the rat genome (Rnor_6.0) and microRNAs were identified using sRNAtoolbox with default settings (6). MicroRNAs were grouped into the classes indicated in Table 1 using the Venn Diagram tool available at http://bioinformatics.psb.ugent.be/webtools/Venn/.

**EV Analysis & *Statistics*.**

The measurements investigating the effect of ES on astrocytes at each frequency were repeated 3 times (Figure 1S). Astrocytes used in each measurement were obtained from independent culture preparations. Significance testing utilized one-way ANOVA and Tukey-Kramer post hoc test (MathWorks, Natick, MA, USA). Each graph of bivariate dot-plots of Alexa488-AQP4 fluorescence (FITC-A) versus side scatter area (SSC-A) in Figure 1S. was split into four regions: **a** (top left, AQP4-carrying exosomes), **b** (top right, AQP4-carrying microvesicles), **c** (bottom left, Non-AQP4-carrying exosomes) and **d** (bottom right, Non-AQP4-carrying microvesicles).  The percentage of the amount of EV in each region was showed at each corner in the graph. The four regions were defined by a FITC-A threshold for AQP4 positive EVs and a EV size of 100 nm to differentiate exosomes and microvesicles. At each frequency the percent change with ES, $\eta_\kappa$, was defined as $\eta_\kappa=\{( V_r'- V_r)/V_r\}_\kappa$,where $V_r$ is the percentage of the amount of EV in each region without ES, $V_r'$ is the percentage of the amount of EV in that region with ES, r is the region name (a, b, c or d) and $\kappa$ is frequency (2 Hz, 20 Hz or 200 Hz). The values of $\eta$ are shown in Table 1S. In Figure 2S the analysis of the 3 independent trials shows that in region **a**, where AQP4-carrying exosomes are, the difference in percent change with ES between 2 Hz, 20 Hz and 200 Hz groups is not statistically significant, $p = 0.530$. In region **b**, where AQP4 carrying microvesicles are, the difference in percent change with ES between 2 Hz, 20 Hz and 200 Hz groups is statistically significant, $p = 0.022$. Tukey-Kramer post hoc analysis showed that the difference in percent change with ES between 2 Hz and 200 Hz is statistically significant, $p = 0.020$. And, there is no statistically significant difference in percent change with ES between 2 Hz and 20 Hz, 20 Hz and 200 Hz.

Figure 1s

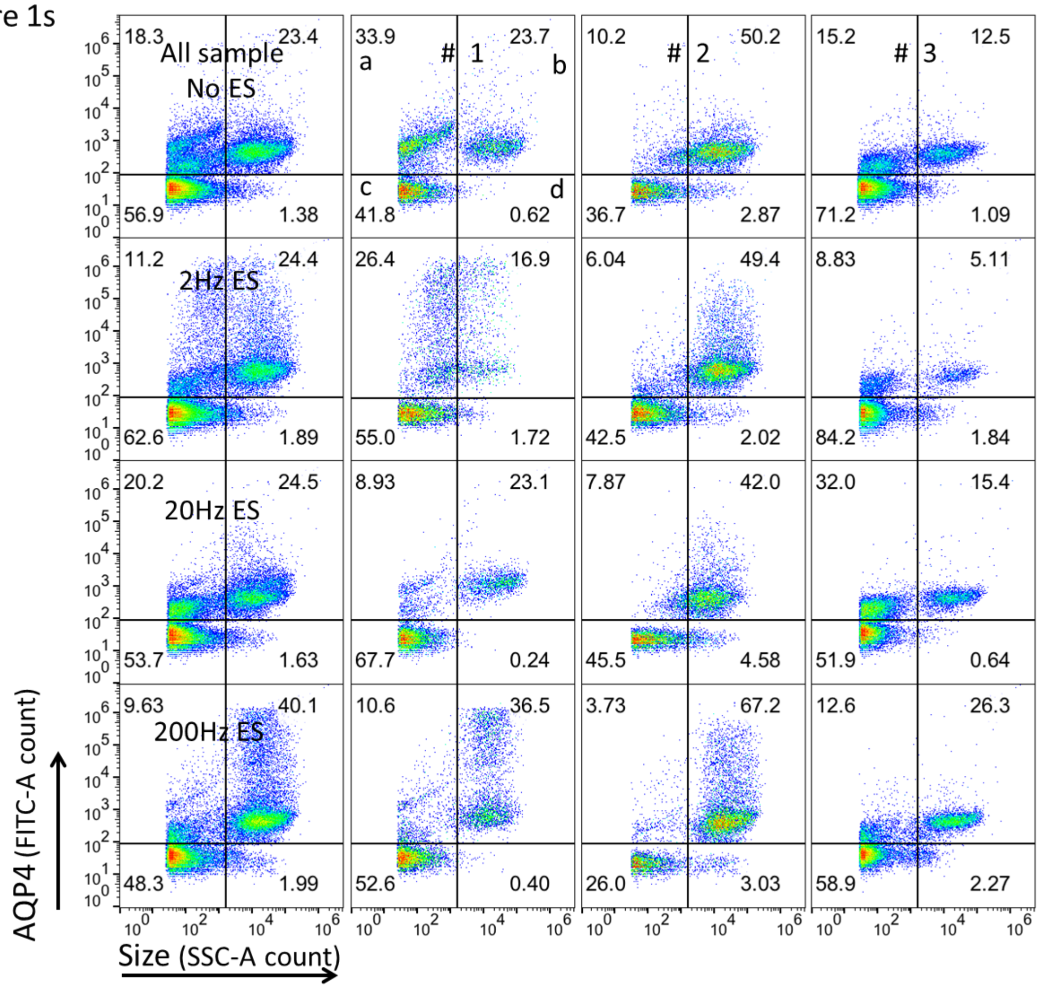

Figure 1S: Three independent trials investigating the effect of electrical stimulation on astrocyte EV release. Left: the aggregated of all 3 trials. Right: the 3 trials

Figure 2s

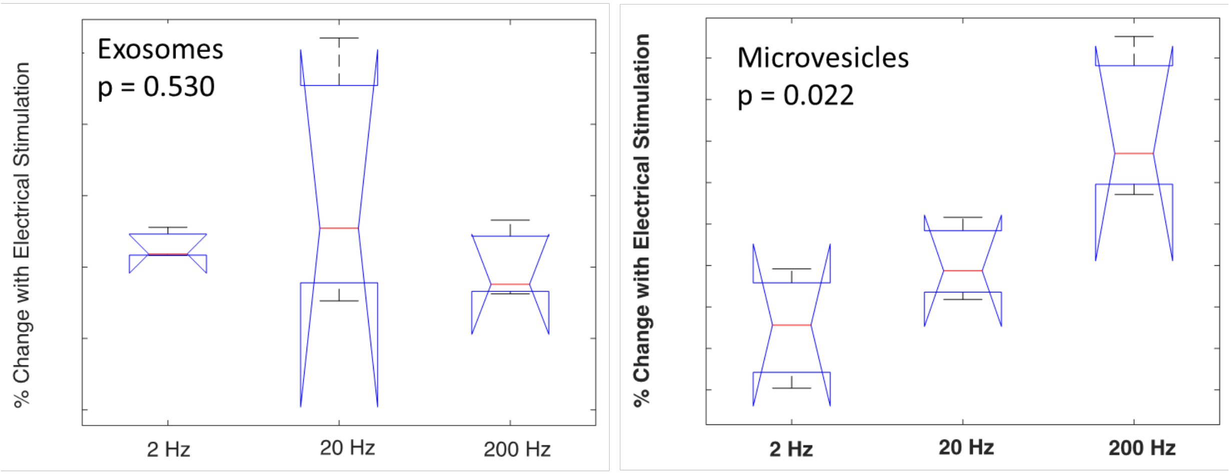

Table 1s

| **Sample_Quad** | **1a** | **1b** | **1c** | **1d** | **2a** | **2b** | **2c** | **2d** | **3a** | **3b** | **3c** | **3d** |
|---|---|---|---|---|---|---|---|---|---|---|---|---|
| **No_ES** | 33.9 | 23.7 | 41.8 | 0.6 | 10.2 | 50.2 | 36.7 | 2.9 | 15.2 | 12.5 | 71.2 | 1.1 |
| **2 Hz** | 26.4 | 16.9 | 55.0 | 1.7 | 6.0 | 49.4 | 42.5 | 2.0 | 8.8 | 5.1 | 84.2 | 1.8 |
| **20 Hz** | 8.9 | 23.1 | 67.7 | 0.2 | 7.9 | 42.0 | 45.5 | 4.6 | 32.0 | 15.4 | 51.9 | 0.6 |
| **200 Hz** | 10.6 | 36.5 | 52.6 | 0.4 | 3.9 | 67.4 | 26.0 | 2.9 | 12.6 | 26.3 | 58.9 | 2.3 |
| **%change, η** | | | | | | | | | | | | |
| **2 Hz** | -0.221 | -0.287 | 0.316 | 1.774 | -0.408 | -0.016 | 0.158 | -0.296 | -0.419 | -0.591 | 0.183 | 0.688 |
| **20 Hz** | -0.737 | -0.025 | 0.620 | -0.613 | -0.228 | -0.163 | 0.240 | 0.596 | 1.105 | 0.232 | -0.271 | -0.413 |
| **200 Hz** | -0.687 | 0.540 | 0.258 | -0.355 | -0.621 | 0.343 | -0.292 | 0.000 | -0.171 | 1.104 | -0.173 | 1.083 |

Figure S2 : ANOVA and Tukey-Kramer post hoc analysis (Matlab). The p is 0.53 for exosomes and 0.022 for microvesicles. The difference in percent change with ES between 2 Hz and 200 Hz is statistically significant ($p = 0.02$) and others are > 0.05.

Table S1. The % change for 2, 20, 200 Hz electrical stimulation in the 3 independent samples shown in Figure S1.

**Supplemental References**

1). McCarthy KD, de Vellis J Preparation of separate astroglial and oligodendroglial cell cultures from rat cerebral tissue. J Cell Biol 1980; 85:890–902.

2). Y. Tian *et al.*, Protein Profiling and Sizing of Extracellular Vesicles from Colorectal Cancer Patients via Flow Cytometry. *ACS nano* 2018; **12**, 671.

3). Zhou K, Spillman MA, Behbakht K, Komatsua JM, Abrahante JE, Hicks DA, Schotl B, Odean E, Jones KL, Graner MW, Bemis LT. A Method for Extracting and Characterizing RNA from Urine: for downstream PCR and RNAseq Analysis. Analytical Biochemistry. 2017; 536:8-15.

4). Seifert VA, Clarke BL, Crossland JP, Bemis LT. A method to distinguish morphologically similar Peromyscus species using extracellular RNA and high-resolution melt analysis. Analytical Biochemistry. 2016:508:65-72.

5). Martin, M. Cutadapt removes adapter sequences from high-throughput sequencing reads. EMBnet Journal 2011; 17:10-11.)

6). Rueda A, Barturen G, Lebron R, Gomez-Martin C, Alganza A, Oliver JL and Hackenberg M. sRNAtoolbox: An integrated collection of small RNA research tools. Nucleic Acids Research. 2015; 43(W1):W467-W4
